# A simple *in vitro* fermentation model to detect alterations in gut microbiota-dependent bile acid profiles

**DOI:** 10.1101/2024.07.31.606071

**Authors:** Weijia Zheng, Wouter Bakker, Marta Baccaro, Maojun Jin, Jing Wang, Ivonne M.C.M. Rietjens

**Author notes:** **Correspondence address:** Weijia Zheng,; Jing Wang.

## Abstract

A previous *in vivo* study showed that lincosamide antibiotics (lincomycin and clindamycin) could induce changes in the gut bacterial community, leading to significant changes in fecal bile acid profiles. Herein, our aim is to develop an animal alternative *in vitro* model for studying gut microbiota-dependent bile acid profiles induced by xenobiotics. The effects of lincosamides were evaluated using this model, and results obtained were verified by comparing with those of the previous *in vivo* study. Fecal sample processing and bile acid incubation conditions were developed and optimized using feces collected from Wistar rats, and prepared samples were incubated for 24 h with or without lincosamides. Upon treatment of the fecal gut microbiota with lincosamides primary and secondary bile acids showed obviously increased and decreased levels respectively. Moreover, the changes in bile acid profile could be linked to a reduced richness of family *Erysipelotrichaceae*, *Bacteroidacea* and *Lactobacillaceae* or *Prevotellaceae*. The consistent consequences of *in vivo* and *in vitro* provides a proof of principle for further application on elucidating effects of other xenobiotics on the gut bacterial community and related bile acid metabolism, thereby contributing to the 3Rs (replacement, reduction and refinement) in animal testing.

## 1. Introduction

It is increasingly appreciated that the gut microbiota contributes to health and disease (Wahlström et al., 2016). It has been reported that the reduced adiposity of germ-free (GF) mice can be reversed by colonization using a normal gut microbiota (Bäckhed et al., 2004); as a result, the gut microbiota has emerged as a critical factor contributing to host health and metabolism. However, the profile of intestinal bacteria could be altered by a variety of factors, especially antibiotics, such as lincosamides (Sung and Lee, 2008). Representatives of lincosamide antibiotics, such as lincomycin and clindamycin, which are normally applied for the treatment of protozoal diseases according to their antibacterial properties (Spížek and Řezanka, 2004).

It is generally believed that the gut microbiota contributes to host metabolism by converting various bioactive compounds, such as bile acids, that may signal to the host by activating cognate receptors in sensitive cells (Holmes et al., 2012). It has been reported that the gut microbiota is involved in the biotransformation of bile acid through deconjugation, dehydroxylation, and reconjugation of these molecules (Hirano et al., 1982). Moreover, bile acids may also display antimicrobial activity that can damage bacterial cell membranes, leading to inhibition of bacterial overgrowth and protection of the liver and intestine against inflammation (Kurdi et al., 2006; Torres-Fuentes et al., 2017). Thus, the altered composition of intestinal microbiota caused by oral administration of antibiotics could further result in the disorder of bile acid profiles and other unknown health effects.

However, bile acid metabolism is complex, since bile acids are initially modified by liver enzymes and upon secretion into the intestines subsequently modulated by the microbiota, which in return regulate the size and composition of the bile acid pool (Forman et al., 1995; Seol et al., 1995). Some studies have reported on antibiotic-induced alterations of gut microbiota and bile acid production in *in vivo* animal models (Behr et al., 2019; Kang et al., 2019; Theriot et al., 2016). Previously, characterization of bile acid pools was mostly achieved from samples of animal *in vivo* work, and thus far, only a few studies have developed *in vitro* batch models to enable quantification of the dynamic intestinal bile acid pool (Martin et al., 2018). In particular, there has been no study reporting on an *in vitro* fermentation model able to mimic and reproduce the alteration of intestinal bile acid profiles observed upon administration of antibiotics or other drugs *in vivo*.

The aim of the present study was to develop and evaluate such an *in vitro* model for studying the potential effects of test compounds on the processing of conjugated bile acids by the rat intestinal microbiota. To this end, a model consisting of anaerobic fecal incubations was characterized in which (altered) bile acid profiles could be quantified using LC‒MS/MS, and bacterial profiles were determined by 16S rRNA gene sequencing analysis. The study design for sample processing and data acquisition of the current study is presented in **Fig. 1**. As a first proof of principle, the *in vitro* model system was applied to study the variation in gut microbiota and bile acid profiles induced by lincosamides to characterize the dynamic interactions between gut microbiota and the bile acid pool and to enable evaluation of the *in vitro* model by comparison to available *in vivo* data on the effect of these antibiotics on bile acid profiles in rats orally exposed to lincomycin and clindamycin (Behr et al., 2019).

**Fig. 1.**
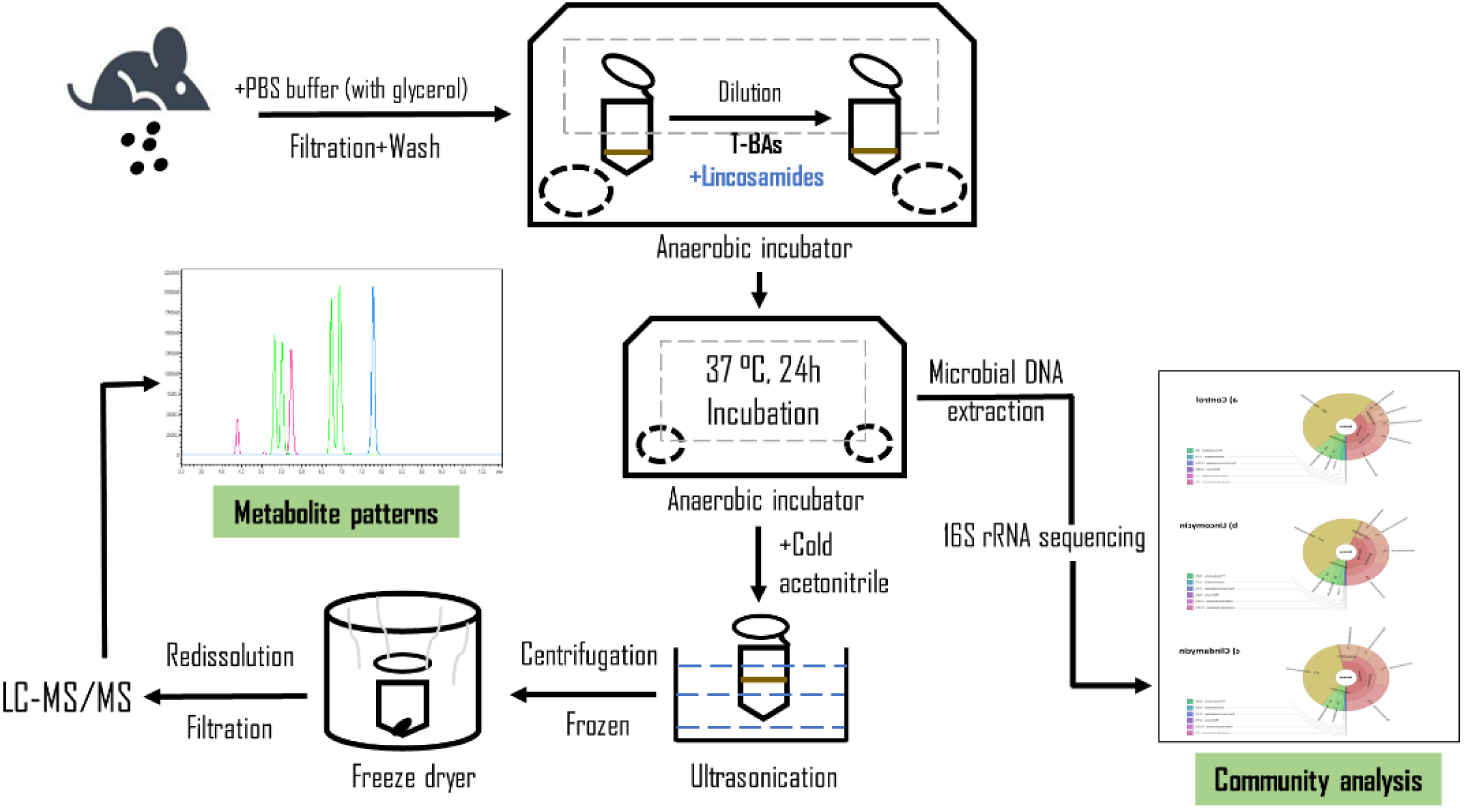
Schematic presentation of the study design. Scheme for bile acid profiling in control and antibiotic-treated rat feces and the corresponding community analysis is shown. Fecal samples for bile acid profiling were taken at 0, 6, 12, and 24 h, while samples for 16S rRNA analysis were taken only at 24 h of anaerobic incubation.

## 2. Material and methods

### 2.1. Chemicals and solvents

The test compounds lincomycin hydrochloride and clindamycin hydrochloride were procured from Sigma Aldrich (Zwijndrecht, The Netherlands). Bile acids were supplied by Merck KGaA (Darmstadt, Germany) or Cambridge Isotope Laboratories (Tewksbury, USA). Methanol and acetonitrile (ACN) were acquired from Biosolve BV (Valkenswaard, The Netherlands), and formic acid was ordered from VWR CHEMICA (Amsterdam, The Netherlands). Phosphate-buffered saline (PBS) was secured from Gibco (Paisley, UK), and para-Pak SpinCon tubes were supplied by Fisher Scientific (New Hampshire, USA). A solution of mixed taurine conjugated bile acids (T-BAs) containing 500 μM taurocholic acid (TCA), 100 μM tauroursodeoxycholic acid (TUDCA), and 100 μM tauro-β-muricholate (TβMCA) was prepared in PBS. The mixture of labelled bile acids ([2,2,4,4-d4] cholic acid and [2,2,4,4-d4] lithocholic acid) obtained from Merck KGaA was dissolved at a final concentration of 100 μM (for each) in methanol as an internal standard for LC‒MS/MS analysis.

### 2.2. Fecal sample preparation

Fresh fecal samples from Wistar rats (20 males and 20 females) were kindly provided by BASF (Ludwigshafen, Germany). Feces from individual animals were obtained by physical stimulation of the abdomen to trigger defecation, after which the fecal samples were weighed and immediately transferred into an anaerobic solution of 10% (*v/v*) glycerol in PBS and diluted to a final fecal concentration of 20% (*w/v*). Pooled samples were manually stirred by a sterile glass wand and flushed with N_2_ gas. Subsequently, samples were filtered using a woven sterile medical gauze dressing (provided by HeltiQ, Wolvega, The Netherlands) under anaerobic conditions (85% N_2_, 10% CO_2_ and 5% H_2_, in a Bactron EZ anaerobic chamber) (Sheldon, Cornelius, USA), and aliquoted samples of the resulting fecal slurry were stored at −80 °C until further use.

### 2.3. Anaerobic incubations and extraction of bile acids

Conditions for the anaerobic incubation of rat feces and subsequent bile acid analysis were optimized to achieve an effective extraction and adequate recovery of the bile acids and to ensure sufficient activities of the gut microbiota. To remove residual endogenous fecal bile acids, filtered fecal samples containing 20% feces (*v/v*) were washed twice using equal volumes of anaerobic PBS (V_fecal sample_: V_PBS_ =1: 1) by vortex mixing for 1 min and centrifugation at 2,000×g for 5 min at 4 °C under anaerobic conditions. Supernatants were removed, and PBS was supplied to obtain the same sample volume as before washing. Subsequently, the samples were incubated with an externally added bile acid mixture. The 200 μL incubation system was composed of 120 μL washed fecal sample (final concentration 1.5 g feces/mL), 20 μL mixed solution of T-BAs providing a final concentration of 50 μM TCA, 10 μM TUDCA and 10 μM TβMCA, 20 μL concentrated stock solutions of lincomycin (final concentration: 16.32 mM) or clindamycin (final concentration: 10.41 mM) in anaerobic PBS buffer or 20 μL PBS for the control without antibiotic. The final concentrations of lincosamides to be tested in the developed *in vitro* model were based on the dose levels used in a literature reported *in vivo* study in rats (Behr et al., 2019) and determined as follows:

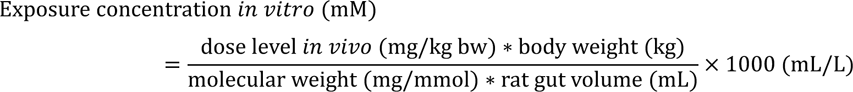

The dose levels used in the reported *in vivo* study amounted to 300 and 200 mg/kg body weight for lincomycin and clindamycin, respectively (Behr et al., 2019), and a rat body weight of 0.25 kg as well as a gut volume of 11.3 mL were applied (Davies and Morris, 1993), as summarized in **Table 1**.

**Table 1.** Calculation of the exposure concentrations of lincosamides used in the fecal incubations.

| Antibiotics | Dose level<br>*in vivo* (mg/kg/bw)<br>(Behr et al., 2018) | Total<br>exposure<br>amount (mg) | Exposure<br>to gut*<br>(mg/mL) | Molecular<br>Weight | Exposure<br>concentration<br>(mM) |
| --- | --- | --- | --- | --- | --- |
| Lincomycin | 300 | 75 | 6.64 | 406.54 | 16.32 |
| Clindamycin | 200 | 50 | 4.42 | 424.98 | 10.41 |

\* A gut volume of 11.3 mL was applied (Davies and Morris, 1993).

Samples were prepared and incubated in an anaerobic chamber (Sheldon, Cornelius, USA) with an atmosphere of 85% N_2_, 10% CO_2_ and 5% H_2_ at 37 °C and terminated at 0 h, 6 h, 12 h and 24 h by adding 200 μL ice-cold acetonitrile to terminate the reaction. Samples were then removed from the anaerobic incubator and placed on ice for 10 min followed by 5 min ultrasonication to rupture the cell membranes of microbes and collect bile acids. Subsequently, centrifugation at 21,500 ×g for 15 min at 4 °C was applied to precipitate microorganisms, particles, and proteins. The upper layers were collected and stored at −80 °C overnight followed by 8-hour freeze drying. The powder obtained was redissolved in 100 μL of nano-pure water and methanol (1/1, *v/v*) and vortexed for 2 min followed by centrifugation at 21,500×g for 5 min. The supernatant was removed and loaded onto a 96-well filter plate (PALL AcroPrep, PTFE 0.2 μm). To 90 μL of filtered extract, 10 µL internal standard was added. The obtained solutions were shaken briefly and subsequently transferred to vials for further analysis by LC‒MS/MS.

### 2.4 Measurement of bile acids by LC‒MS/MS

LC–MS/MS detection and quantification of bile acids was performed using a Shimadzu Nexera XR LC-20CE SR UPLC system coupled with a Shimadzu LCMS-8050 mass spectrometer (Kyoto, Japan) with an electrospray ionization (ESI) interface. Electrospray ionization (ESI) in turbo negative ion mode was performed with the triple quadrupole tandem mass spectrometric (MS/MS) system, and multiple reaction monitoring was utilized for data acquisition. Individual standard solutions (1 μg/mL) of each of the bile acids were employed to optimize the precursor ion, product ion, declustering potential and collision energy by direct injection into the mass spectrometer. The chemical structures, as well as parent and fragment masses of 24 major unconjugated bile acids and their glycine and taurine conjugates, are shown in **Table 2**. Samples were loaded, and the target bile acids were separated on a Kinetex 100A C18 column (2.1*50 mm, 1.7 mm) provided by Phenomenex (Torrance, USA). The mobile phases consisted of 0.01% formic acid in distilled water (solvent A), a mixture of methanol and acetonitrile (v/v=1/1) (solvent B), and acetonitrile containing 0.1% formic acid (solvent C). The column temperature was 40 °C, the flow rate was 0.5 mL/min, and the injection volume was 2 μL. The total run time was 16 minutes with the following gradient profile: 95% A, 0% B and 5% C (0-2 min), slowly changed to 30% A, 70% B, and 0% C from 2 to 7.5 min, rapidly reversed to 2% A, 98% B, and 0% C in 0.1 min, then kept at 2% A, 98% B, and 0% C from 7.6 to 10 min, then changed to 70% A, 30% B, and 0% C in 0.5 min; in the end, slowly returned to the initial conditions of 95% A, 0% B and 5% C from 10.5 to 13 min, then maintained at these conditions until 16 min for equilibration.

**Table 2.** MRM data acquisition parameters of the LC‒MS/MS procedure for unconjugated bile acids and conjugated bile acids. Unconjugated bile acids involve primary bile acids (CA, CDCA, αMCA and βMCA) and secondary bile acids (DCA, LCA, UDCA, HDCA, and ωMCA). Conjugated bile acids include tauro-bile acids (X=taurine) and glycol-bile acids (X=glycine).

| Analytes | Precursor<br>Ion (*m/z*) | Product<br>Ion (*m/z*) | Collision<br>energy (eV) | R1 | R2 | R3 | X |
| --- | --- | --- | --- | --- | --- | --- | --- |
| Cholic acid (CA) | 407.3 | 407.3 | 25 | -H | -OH(α) | -OH | -OH |
| Taurocholic acid (TCA) | 514.4 | 514.4 | 30 | -H | -OH(α) | -OH | -taurine |
| Glycocholic acid (GCA) | 464.3 | 74.0 | 43 | -H | -OH(α) | -OH | -glycine |
| Chenodeoxycholic acid (CDCA) | 391.3 | 391.3 | 15 | -H | -OH(α) | -H | -OH |
| Taurochenodeoxycholic acid (TCDCA) | 498.4 | 498.4 | 55 | -H | -OH(α) | -H | -taurine |
| Glycochenodeoxycholic acid (GCDCA) | 448.3 | 74.0 | 43 | -H | -OH(α) | -H | -glycine |
| Deoxycholic acid (DCA) | 391.3 | 391.3 | 15 | -H | -H | -OH | -OH |
| Taurodeoxycholic acid (TDCA) | 498.4 | 498.4 | 55 | -H | -H | -OH | -taurine |
| Glycodeoxycholic acid (GDCA) | 448.3 | 74.0 | 43 | -H | -H | -OH | -glycine |
| Lithocholic acid (LCA) | 375.3 | 375.3 | 30 | -H | -H | -H | -OH |
| Taurolithocholic acid (TLCA) | 482.3 | 482.3 | 54 | -H | -H | -H | -taurine |
| Glycolithocholic acid (GLCA) | 432.3 | 74 | 55 | -H | -H | -H | -glycine |
| Ursodeoxycholic acid (UDCA) | 391.3 | 391.3 | 15 | -H | -OH(β) | -H | -OH |
| Tauroursodeoxycholic acid (TUDCA) | 498.4 | 498.4 | 55 | -H | -OH(β) | -H | -taurine |
| Glycoursodeoxycholic acid (GUDCA) | 448.3 | 74.0 | 43 | -H | -OH(β) | -H | -glycine |
| Hyodeoxycholic acid (HDCA) | 391.3 | 391.3 | 15 | -OH(α) | -H | -H | -OH |
| Taurohyodeoxycholic acid (THDCA) | 498.4 | 498.4 | 55 | -OH(α) | -H | -H | -taurine |
| Glycohyodeoxycholic acid (GHDCA) | 448.3 | 74.0 | 43 | -OH(α) | -H | -H | -glycine |
| α-Muricholate (αMCA) | 407.3 | 407.3 | 25 | -OH(β) | -OH(α) | -H | -OH |
| Tauro-α-muricholate (TαMCA) | 514.4 | 514.4 | 30 | -OH(β) | -OH(α) | -H | -taurine |
| β-Muricholate (βMCA) | 407.3 | 407.3 | 25 | -OH(β) | -OH(β) | -H | -OH |
| Tauro-β-muricholate (TβMCA) | 514.4 | 514.4 | 30 | -OH(β) | -OH(β) | -H | -taurine |
| ω-Muricholate (ωMCA) | 407.3 | 407.3 | 25 | -OH(α) | -OH(β) | -H | -OH |
| Tauro-ω-muricholate (TωMCA) | 514.4 | 514.4 | 30 | -OH(α) | -OH(β) | -H | -taurine |

#### Microbial Taxonomic Profiling and Total Bacterial Load

A total of 240 µL diluted and filtered fecal slurry containing 20% feces (*v/v*) (as described in the “fecal sample preparation” section) and 30 µL PBS mixed with 30 µL stock solution of lincomycin (final concentration: 16.32 mM) or clindamycin (final concentration: 10.41 mM) was incubated under anaerobic conditions for 24 h at 37 °C, and stored at −80 °C overnight (3 aliquots per group). Subsequently, DNA was isolated from the fecal slurries using a bead-beating procedure coupled with the customized MaxWell 16 Tissue LEV Total RNA Purification Kit (XAS1220; Promega Biotech AB, Stockholm, Sweden). DNA isolates underwent triplicate polymerase chain reaction (PCR) reactions of the 16S rRNA gene V4 region (515-F; 806-R), and PCR products acquired were purified, pooled, and sequenced (Illumina NovaSeq 6000, paired-end, 70 bp; Eurofins Genomics Europe Sequencing GmbH, Konstanz, Germany).

To quantify the total bacterial load in each individual fecal slurry, quantitative PCR (qPCR) was carried out. PCR products were generated by amplification using 16S V3−V4 primers (F-NXT-Bakt-341F: 5′-CCTACGGGNGGCWGCAG-3′ and R-NXT-Bakt-805R: 5′-GACTACHVGGGTATCTAATCC-3′). During index PCR, barcodes for multiplexed sequencing were introduced using overhang tags. A sequencing library was prepared from the barcoded PCR products and sequenced on the Illumina MiSeq next-generation sequencing system (Illumina Inc.). Signals were processed to *.fastq-files, and the resulting 2 × 250 bp reads were demultiplexed. Microbiota identification was performed by clustering the sequences at a 97% identity threshold defining operational taxonomic units (OTUs), according to the taxonomy of the SILVA 132 16S rRNA sequence database.

#### Data Analysis

Metabolic profile data acquisition and processing were carried out using the Labsolutions software of the LC-TQ-MS system. Graphics were presented by using GraphPad Prism 8.2 (San Diego, USA), and chemical structures were drawn by using ChemDraw 18.0 (PerkinElmer, Waltham, USA). The results are shown as the mean ± standard deviation (SD).

## 3. Results and discussion

### Bile acid measurement by LC‒MS/MS

**Fig. 2 A-C** show the chromatograms of unconjugated bile acids, glycine conjugated bile acids and taurine conjugated bile acids, respectively. Good separation with an identified retention time for each bile acid and high sensitivity with limits of quantification ≤0.05 μM for all targets were obtained after the development and optimization of the LC‒MS/MS detection method.

**Fig. 2.**
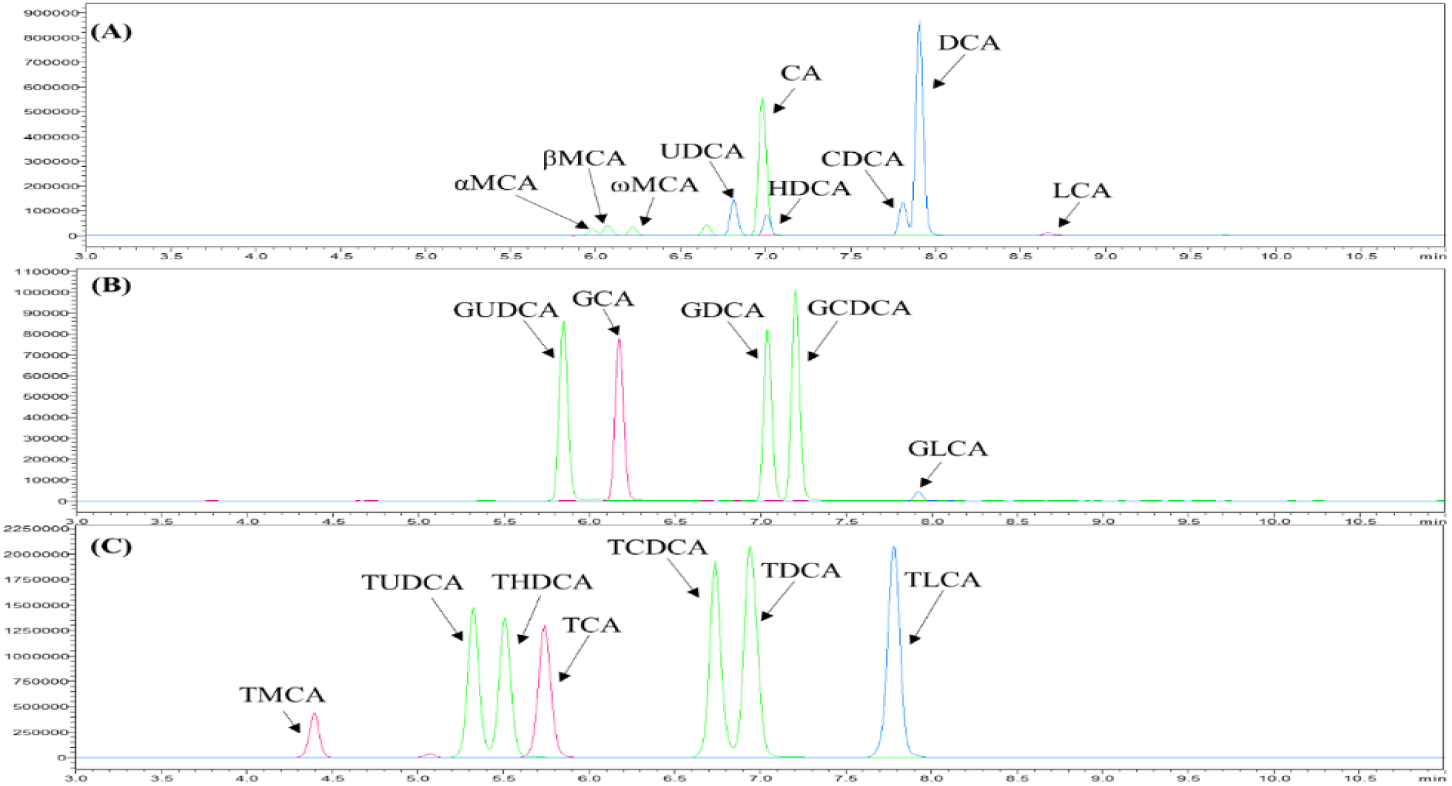
Chromatography. UPLC chromatography graphs of (A) unconjugated bile acids (blue: 391.3>391.3; green: 407.3>407.3), (B) glycine conjugated bile acids (blue: 432.3>74.0; green: 448.3>74.0; pink: 464.3>74.0) and (C) taurine conjugated bile acids (blue: 482.3>482.3; green: 498.4>498.4; pink: 514.4>514.4) in a mixed standard solution with final concentration of 1 μM standard for each. The time range displayed is from 3 to 11 min.

To obtain a higher intensity of signals for all these bile acids, a pseudo-MRM method, which applies the same product ion as the parent ion under an optimized collision energy, was employed. Since all these unconjugated bile acids have no 12-hydroxyl group that is fragmented merely via dehydration/dehydrogenation, no discriminative fragments were exhibited regardless of sites and epimerization of the hydroxyl groups on the skeleton (Lan et al., 2016). Thus, fragments such as *m/z* 407.3 > 407.3 for CA and *m/z* 391.3 > 391.3 for CDCA were used for detection of the unconjugated isomers and stereoisomers (Yin et al., 2017). Upon testing both ESI^+^ and ESI^-^, the negative ionization mode was selected since it provided more stable and stronger signals. **Table 2** summarizes the MRM data acquisition parameters of the LC‒MS/MS procedure for unconjugated bile acids and conjugated bile acids.

To optimize the chromatographic separation of the bile acids, various mobile phases with various counterions in the pH range of 3∼10 were evaluated, since due to their different pKa values, different effects of acidity on the retention of bile acids were observed. The unconjugated bile acids and glycine conjugates usually have pKa values in the range of 4∼6, while the pKa values of taurine conjugates are less than 2.0 (Carey et al., 1972). Therefore, decreasing the pH of the mobile phases markedly increased the retention of glycine-conjugated bile acids and unconjugated bile acids, which improved their chromatographic separation with minimal influence on taurine bile acids. Meanwhile, the ionization efficiency and therefore the signal intensity in ESI^-^ mode are also strongly dependent on mobile phase constituents and pH (Alnouti et al., 2008). Thus, based on our findings, the mobile phases separately adding 0.01% and 0.1% formic acid were finally selected in a gradient of methanol, acetonitrile, and distilled water.

In addition, the hydrophilicity of these bile acids based on their nucleus and side chain structures is another important factor influencing chromatographic separation. For example, tri-hydroxy bile acids (CA, MCA) elute earlier than di-hydroxy bile acids (CDCA and DCA), which in turn elute earlier than mono-hydroxy bile acids (LCA). This elution behaviour may be attributed to the orientation of the hydroxyl substitutions and their ability to form intramolecular H-bonds (Ikegawa et al., 1996). According to this, a reversed-phase C18 column was tested and selected to obtain the optimized separation for all the targets.

### Development and optimization of the *in vitro* anaerobic fecal incubation model

An anaerobic fecal batch culture model in PBS was developed to study bacterial metabolism. **Fig. 3** shows the bile acid levels in incubations with freshly isolated (no treatment), PBS-washed, and T-BA supplemented fecal samples at t=0, is the latter also reflecting the ultimate optimized protocol applied for subsequent studies on the effects of the selected antibiotics (**Fig. 3C**). In samples with freshly isolated rat feces, both secondary bile acids (ωMCA, HDCA, DCA and LCA) and primary bile acids (βMCA, αMCA, and CA) were readily detected, as shown in **Fig. 3A**. Glycine-conjugated bile acids (G-BAs) were not detected, which is in line with a previous study reporting a low proportion of G-BAs (<1%) in rat liver (Lin et al., 2019). Levels of T-BAs were present below the detection limits, while these are known to be readily excreted from the liver into the intestines. This implies that the bile acid profile in freshly isolated fecal samples does not reflect the bile acid profile in the intestine before metabolism by the gut microbiota as such. To define a bile acid content of the fecal incubations more in line with what would be expected in the intestines, fecal samples were washed and supplemented with a mixture of T-BAs. After washing the freshly isolated rat fecal samples with PBS, the amount of the detected bile acids was significantly reduced, leaving ωMCA, HDCA, βMCA and DCA at levels amounting to 8.58%, 12.67%, 9.74%, and 25.84% of the levels detected in the freshly isolated fecal samples (**Fig. 3**). Subsequently, to obtain the final incubations, TCA, TβMCA, and TUDCA were added to the fecal incubations at levels amounting to final target concentrations of 50 μM TCA, 10 μM TUDCA, and 10 μM TβMCA to better simulate the bile acid composition of the gut environment. The levels and total amount of TCA, TβMCA, and TUDCA were chosen based on a previous study (Lin et al., 2019), which reported a composition of 70.82% TCA, 13.47% TβMCA, and 11.08% TUDCA present in rat liver and thus expected to be transferred to the intestinal lumen. Thus, a rough ratio of 5:1:1 for TCA, TβMCA, and TUDCA was added to imitate the *in vivo* gut levels of these bile acids. Furthermore, as a result of the apparently extremely fast deconjugation of these T-BAs, at t=0, the incubations contained the conjugated bile acids TCA, TUDCA, TβMCA, and TαMCA at concentrations somewhat lower than the added amounts while also substantial levels of the corresponding deconjugated primary bile acids were detectable (**Fig. 3** and **Table S1**). For example, upon supplementation of the washed samples with TCA, CA instantly increased as well accompanied by an instant reduction of the TCA levels from 50 mM (added) to approximately 30 mM (detected), pointing at apparently swift deconjugation of TCA. In this way, fecal incubations with a reasonable combination of conjugated, primary and secondary bile acids could be obtained, reflecting intestinal levels better than the residual bile acid levels in the fecal samples.

**Fig. 3.**
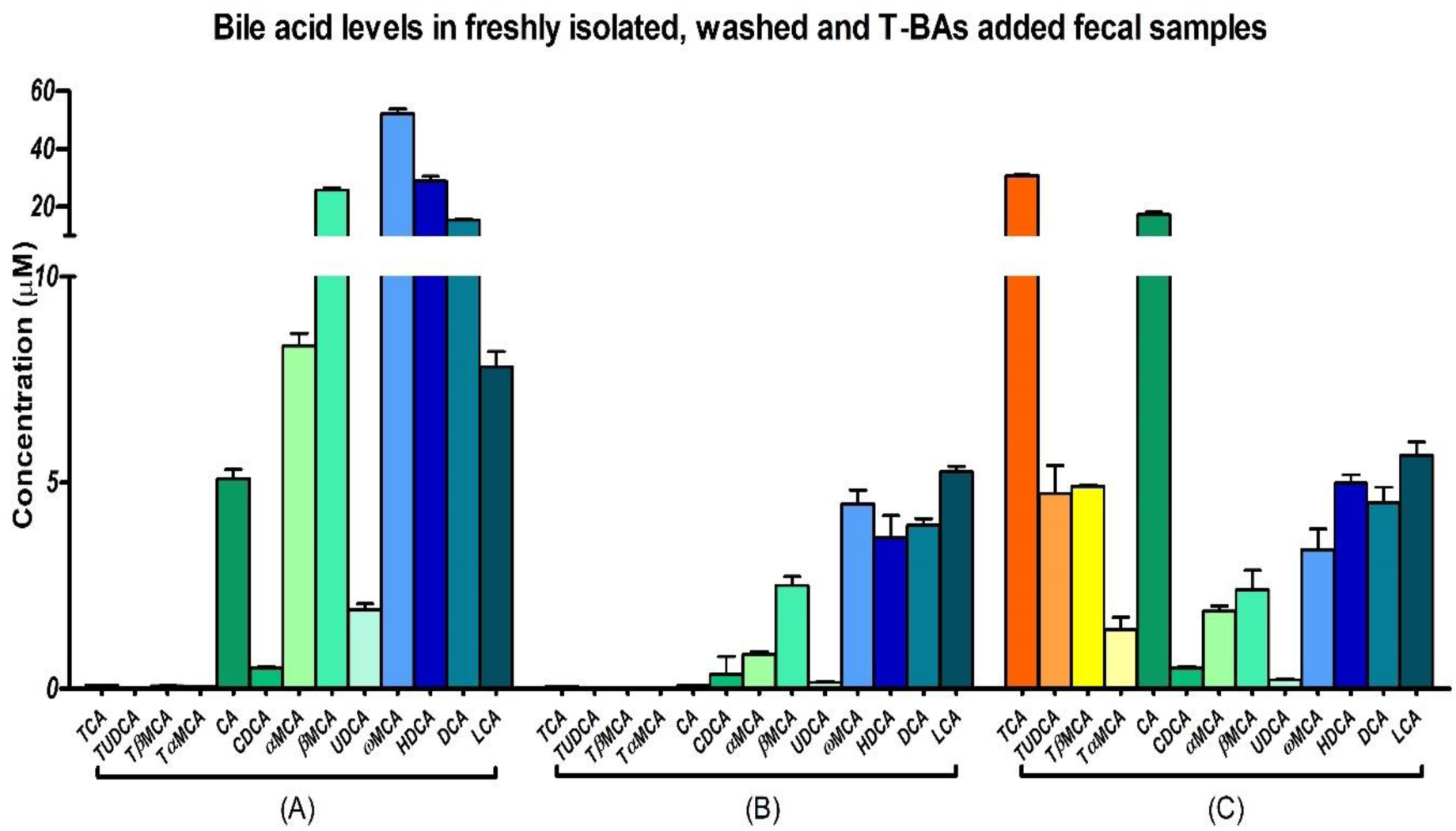
Bile acid profiles. Bile acid pool at t=0 h in incubations with 20% rat feces (v/v) from (A) freshly isolated rat fecal samples, (B) freshly isolated rat fecal samples washed twice using anaerobic PBS, and (C) washed fecal samples with the addition of a mixed T-BA solution of TCA, TUDCA, and TβMCA at final concentrations of 50 μM, 10 μM, and 10 μM Data presented are the mean ± SD from 3 independent experiments.

It is also of importance to note that some bile acids, such as DCA, LCA, CDCA and their derivatives, have strong hydrophobicity (Májer et al., 2014), and high concentrations could alter bacterial membrane permeability, leading to disturbed gut microbiota (Islam et al., 2011; Ramírez-Pérez et. al, 2018). Thus, the modification of the initial bile acid pool at the beginning of the incubation was also used to avoid such a negative effect on bacteria. A washing step using anaerobic PBS (equal volume as the filtered fecal sample) followed by centrifugation at 2,000×g for 5 min was applied. Following these optimization steps, the total amount of T-BAs and unconjugated bile acids present in the incubations at t=0 were comparable at the start of the incubations (**Table S1**). The supplemented sample thus obtained was used for further studies on the effects of antibiotics on gut microbiota and the resulting effects on bile acid composition and metabolism.

In addition to the modification and optimization of the initial bile acid profile in the *in vitro* fecal incubations, the time of incubation was also optimized. This is because the variation in bacteria upon prolonged incubation times could affect the reproducibility and linearity of the results. In a previous study on conjugated bile acid processing by the human gut microbiota, an incubation period of 24 h was applied, which is relevant for intestinal transit times (Martin et al., 2018); finally, samples taken at 0, 6, 12, and 24 h were selected for final bile acid analysis. Additionally, ice-cold acetonitrile was utilized for not only stopping the reactions but also as the solvent to extract the bile acids efficiently from the fecal incubations.

### Alteration of bile acids

After optimization of the conditions for the anaerobic fecal incubations and the subsequent sample preparation protocol, incubations in the absence and presence of the selected antibiotics lincomycin and clindamycin were performed. **Fig. 4 A** presents the changes in bile acid composition during 24 h anaerobic incubations in the absence (control) or presence of lincomycin and clindamycin. From these data, it follows that during the 24 h lincosamide treatment of the fecal samples, resulted in significant changes in the bile acid profile. **Fig. 4 B** shows the principal component analysis (PCA) according to the bile acid profiles in the antibiotic-treated and control samples taken at 24 h of incubations, and the clustered controls with an obvious separation with lincomycin and clindamycin treated groups reveals the effects of lincosamides on gut-mediated bile acids metabolism.

**Fig. 4.**
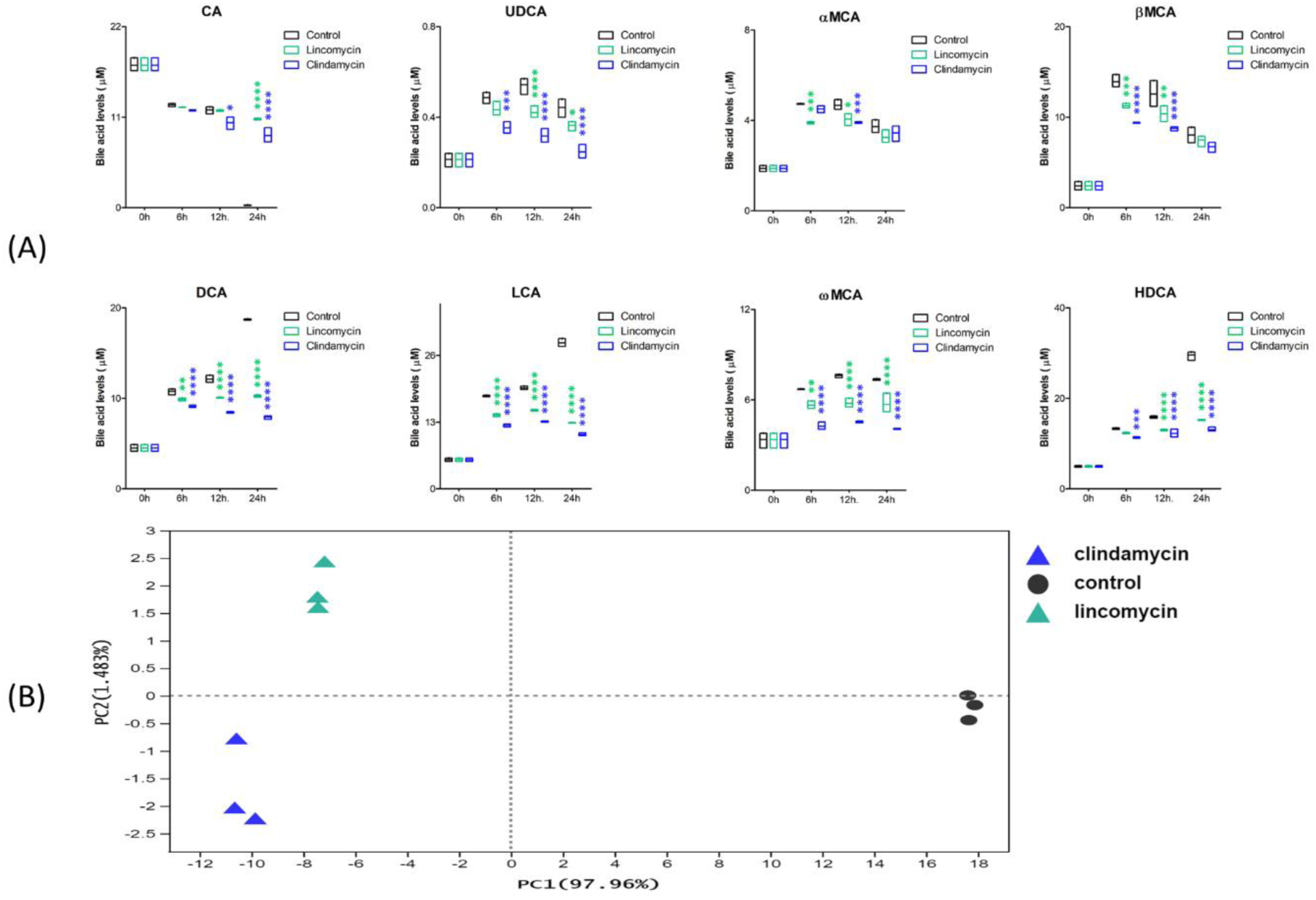
Altered bile acid profiling. Time-dependent changes in bile acid concentrations in anaerobic fecal incubations at 0, 6, 12, and 24 h in control (black), lincomycin-treated (green), and clindamycin-treated (blue) fecal samples are shown in (**A**) (n=3 per group). The bile acids of CA, αMCA, βMCA, UDCA, DCA, LCA, ωMCA, and HDCA were detected and quantified in control and treated samples taken from the anaerobic fecal incubations at the time points indicated (****p <0.0001, *** 0.0001≤ p <0.001, ** 0.001≤ p <0.01, * 0.01≤ p <0.05 indicates a difference of the antibiotic treated samples from the control samples without antibiotic at the corresponding time point). The principal coordinate analysis (PCoA) based on the bile acid profiles of control and treated fecal samples taken at 24 h is shown in (**B**).

In line with the optimized conditions (**Fig. 3 C**) at the start of the incubations, especially the primary bile acids CA, αMCA, and βMCA (except for CDCA), and the secondary bile acids DCA, LCA, ωMCA, and HDCA were detected along with a minor amount of UDCA. In the samples treated with either lincomycin or clindamycin, effects on bile acid pools were readily observed compared to the control incubation, with changes being most pronounced upon 24 h of incubation. During the 24 h incubation, the levels of most bile acids changed to a significantly larger extent in the presence of the antibiotics than in the control, in most cases resulting in lower concentrations, while for CA, the opposite was observed, resulting in higher levels of CA at the end of the 24 h incubation. The CA levels in the control were as low as 0.3±0.04 μM at the end of the 24 h incubation, while 50% of the original amount of CA was still detected in the lincomycin- and clindamycin-treated samples. These higher residual levels of CA in both antibiotic-treated samples were accompanied by a comparable reduction in the level of its secondary bile acid DCA, indicating an effect of the antibiotics on the bacteria responsible for this conversion. Similar effects were observed for the formation of other secondary bile acids, including UDCA, LCA, and HDCA. In particular, the production of DCA, LCA and HDCA in the antibiotic-treated samples was only half of that in the control group. The effects of the antibiotics on αMCA and βMCA were less pronounced and not significant at the end of the 24 h incubation. In line with the rapid deconjugation observed when optimizing the incubation protocols, the taurine-conjugated bile acids TCA, TUDCA, TαMCA, and TβMCA, which were present at the start of the incubations at levels of approximately 42 mM in total (**Table S1**), were almost completely deconjugated at the first time point in all incubations.

### Evaluation of the *in vitro* model for studying alterations in microbiota-dependent bile acid metabolism by comparison to *in vivo* data

To evaluate the use of the *in vitro* incubations for elucidating effects on microbiota-mediated bile acid metabolism, **Fig. 5** presents a comparison of the changes observed in the bile acid profiles induced by lincomycin and clindamycin in the newly defined *in vitro* model and data reported in a previous *in vivo* study (Behr et al., 2019). The overview thus obtained reveals that for most of the primary and secondary bile acids studied, including CA, βMCA, DCA, LCA, ωMCA, and HDCA, the effects observed in this *in vitro* model match the *in vivo* data, with increased levels of CA and decreased or unmodified levels for the other bile acids upon antibiotic treatment. For the remaining two bile acids, UDCA and αMCA, the *in vivo* data in males and female rats did not match, and the female data are in line with what was observed in the *in vitro* model using mixed gender fecal samples. Taken together, it is concluded that upon the treatment of the gut microbiota with antibiotics, similar perturbations of gut microbiota-mediated bile acid metabolism were found in the *in vitro* model and the *in vivo* situation for both primary and secondary bile acids. In addition, **Fig. 6** shows the bioconversion pathways of added TCA, TβMCA, and TUDCA to further explain the alteration of the bile acid profile.

**Fig. 5.**
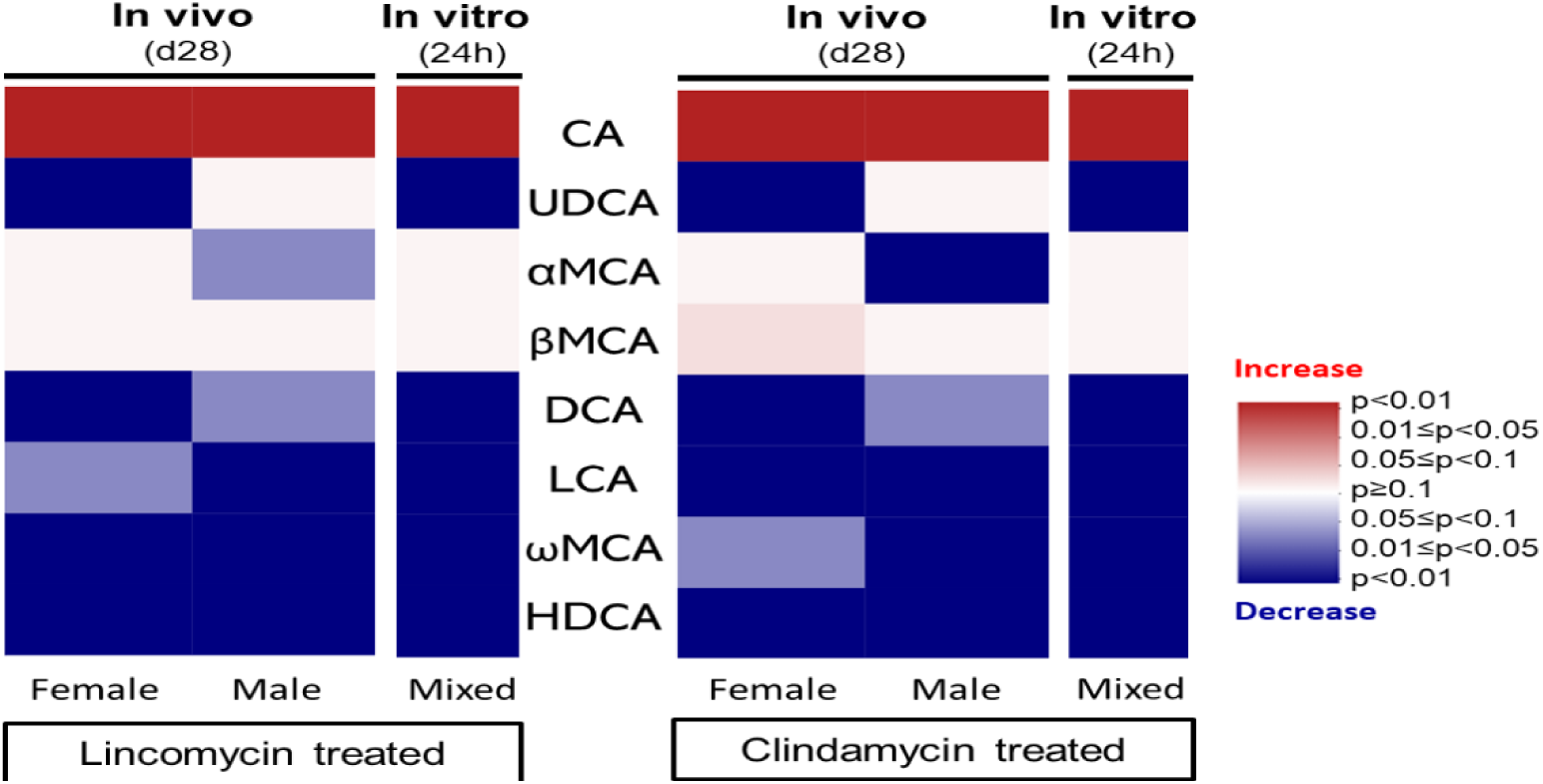
Comparison of *in vitro* and in *vivo*. Comparison of the effect of lincomycin and clindamycin on bile acid concentrations in the *in vitro* anaerobic fecal incubations (mixed gender) at 24 h with the *in vivo* effects detected in feces of male and female rats exposed to lincosamides vs. feces of control rats on day 28 (Behr et al., 2019). Red boxes indicate a significant increase, blue boxes indicate a significant decrease, and the intensity of the color corresponds to the significance of the change. P values are presented in light and dark colors.

**Fig. 6.**
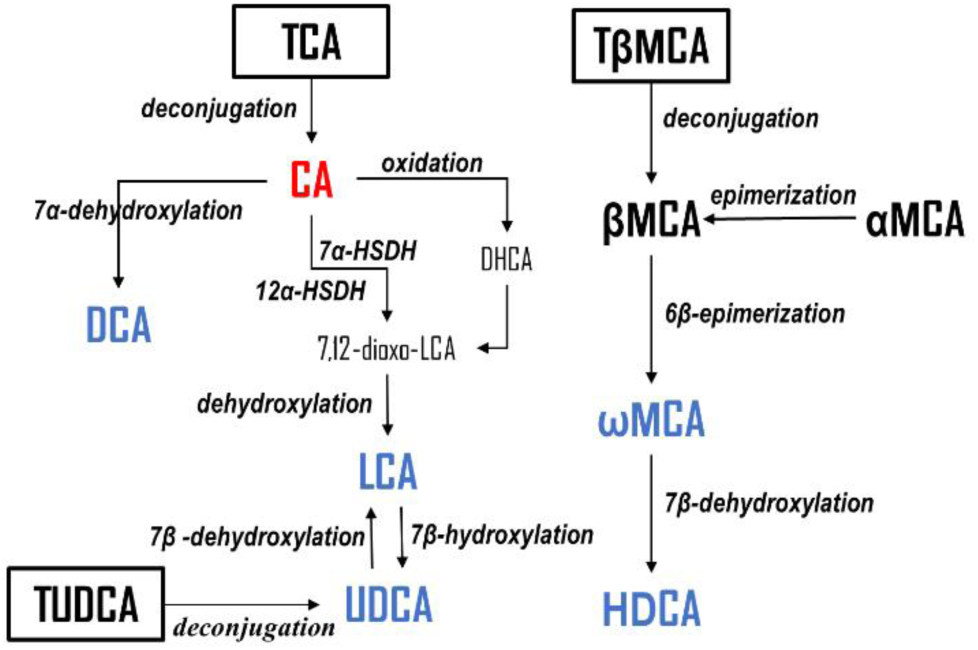
Pathways of fecal T-BA transformation. T-BAs (in black boxes), including TCA, TβMBA, and TUDCA, transform to CA, βMCA, and UDCA by deconjugation; then, the primary bile acids CA and βMCA (some of which can be produced from αMCA) are further converted to various secondary bile acids, including DCA, LCA, UDCA, ωMCA, and HDCA, through 7α/β-dehydroxylation, 7β-hydroxylation, epimerization, and oxidation (Zhang and Klaassen, 2010; Wahlström et al, 2016). In addition, on the pathway of CA transforming to LCA, the intermediate 7,12-dioxo-LCA can be generated via a special oxidation by 7α-hydroxysteroid dehydrogenase (7α-HSDH) and 12α-hydroxysteroid dehydrogenase (12α-HSDH) produced by gut microbiota (Tonin and Arends, 2018). Bile acids showing a significant increase or decrease in the present study are colored red and blue, respectively.

The results of both the *in vivo* (Behr et al., 2019) and the present *in vitro* study reveal the inhibition of CA decomposition upon antibiotic treatment, and these findings have also been reported in other *in vivo* studies (Shimizu et al., 2011; Theriot et al., 2016). This increase in levels of CA and the accompanying decrease in DCA as the primary product of CA upon treatment of the microbiota with antibiotics, observed in both the current *in vitro* and the literature reported *in vivo* study (Behr et al., 2019), indicate a reduction in the 7α-dehydroxylation activity of the intestinal microorganisms (shown in **Fig. 6**). In addition to the formation of the major CA metabolite DCA, **Fig. 6** also shows that a small amount of CA can be converted to the secondary bile acid LCA via oxidation and dehydroxylation. The accumulated CA and decreased production of LCA revealed a slower oxidation/dihydroxylation induced by antibiotic administration as well. Notably, LCA and UDCA can be converted to each other by 7β-hydroxylation and 7β-dehydroxylation, respectively (Wahlström et al, 2016; Tonin and Arends, 2018); therefore, the reduction in LCA could be accompanied by a reduction in UDCA.

On the other hand, the added TβMCA rapidly deconjugated to βMCA, followed by conversion to ωMCA and further transformation to HDCA (**Fig. 6**). Additionally, it was observed (**Fig. 5**) that the production of ωMCA and HDCA was significantly reduced upon treatment with the antibiotics, while an alteration of βMCA was not obvious. Therefore, the slower production of ωMCA could be linked to weakened 6β-epimerization, leading to a reduced level of HDCA. Meanwhile, the βMCA levels may be dependent on a dynamic equilibrium between weakened 6β-epimerization and stronger epimerization of αMCA, the latter in line with the decreased level of αMCA in antibiotic-treated male rats (Wahlström et al, 2016), providing support for this speculation. In addition, sex differences were found in the *in vivo* study with respect to the levels of UDCA, αMCA, and βMCA (only in the clindamycin-treated group), as reported in some previous studies where obvious differences were found between male and female mice in metagenomic analysis involving bile acid metabolism (Sheng et al., 2017; Xie et al., 2017).

### Microbial Taxonomic

**Fig. 7A** and **B** presents the results of the 16S rRNA analysis of the gut microbiota composition of the fecal samples from control rats incubated *in vitro* either without (control) or with lincomycin or clindamycin, showing the relative microbial profile for the phylum and dominant families. The figures reveal that the altered abundance of bacterial species is relatively small but consistent. At the phylum level, the *in vitro* fecal microbial community upon 24 h of *in vitro* incubation for the control sample was composed mainly of *Firmicutes* and *Verrucomicrobia*, followed by *Bacteroidetes* and *Proteobacteria*. In lincomycin- or clindamycin-treated samples, *Proteobacteria* abundance was obviously reduced compared to that in the control group, whereas the relative abundance of *Verrucomicrobia* and *Bacteroidetes* showed a slight increase and decrease, respectively. At the family level (**Fig. 7B**), *Akkermansiaceae*, *Ruminococcaceae*, and *Lachnospiraceae* dominated in rat fecal microbial communities, and no obvious alterations of the three bacterial species were shown in the antibiotic-treated samples compared to controls. In converse, the relative abundance of *Erysipelotrichaceae* significantly reduced in both antibiotic-treated samples (0.0001< p <0.001), and the richness of *Bacteroidaceae* also markedly reduced in lincomycin (0.01≤ p <0.05) and clindamycin (0.0001<p<0.001) groups compared to control. In addition to these changes, the family of *Prevotellaceae* and *Lactobacillaceae* also presented a significant alteration but differently in the antibiotic-treated groups. Relative abundance of *Prevotellaceae* showed the obviously decrease only in the clindamycin-treated fecal samples (0.001≤ p <0.01), while the richness of *Lactobacillaceae* only significantly reduced in the lincomycin-treated group (0.01≤ p <0.05). Although the discrepancy of changes was shown between the bacterial profiles in different antibiotic-treated samples (**Fig. 7B**), effects of the antibiotics on the rat gut microbiome were obviously revealed and further described by the principal coordinate analysis (PCoA) of the Bray‒Curtis distance matrix based on the obtained bacterial profiles (OTUs) in **Fig. 7C**, showing the clustered controls with a significant separation from the antibiotic-treated fecal samples. Moreover, the alpha diversity of the samples with or without *in vitro* antibiotic treatment presented by the Chao and Shannon indexes was shown in **Fig. 7D**, revealing the biodiversity and the relative abundance of species. The total amount as well as biodiversity of the bacterial species presented a slight increase and decrease respectively in the lincomycin and clindamycin treated groups, with no significance shown.

**Fig. 7.**
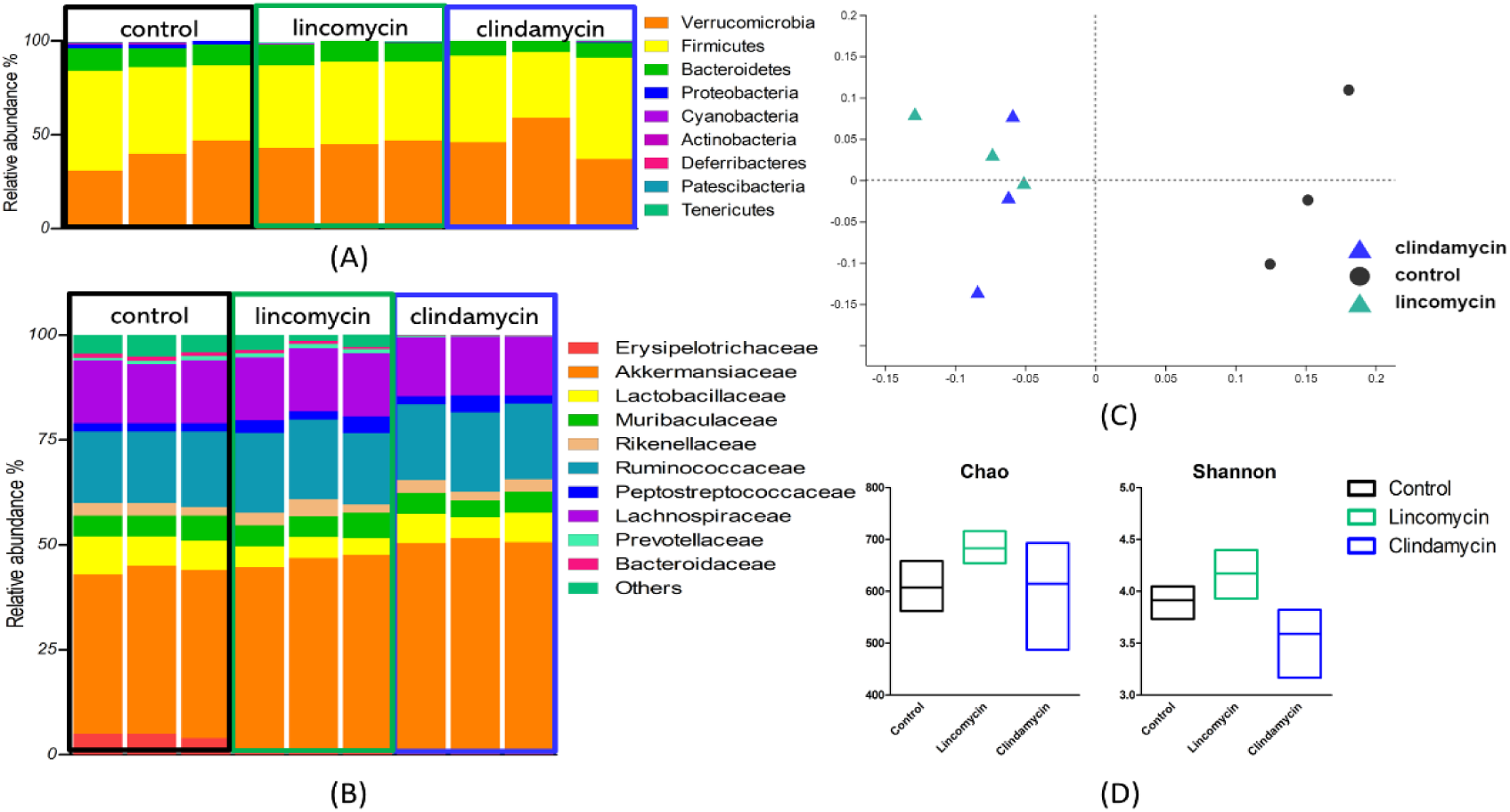
Altered bacterial profiles. Relative abundance of gut microbes in control and lincomycin-or clindamycin-treated pooled fecal samples (n=3) from Wistar rats at the (A) phylum and (B) dominant family levels are described and all the samples were taken after a 24 h incubation. Based on the data of OTUs obtained from these samples, the principal coordinate analysis (PCoA) of the Bray‒Curtis distance matrix and the alpha diversity measured by the Chao and Shannon index are shown in (C) and (D) respectively, plotted for control (black), lincomycin-treated (green), and clindamycin-treated (blue) fecal samples.

The studies published previously which focused on the gut microbial community induced by the exposure of antibiotics achieved the same results as those observed in the present study. For example, a previous mouse *in vivo* study has reported the changes in gut microbial communities induced by gentamicin, and *Erysipelotrichaceae* was also found to be mutated further leading to changes in short-chain fatty acids and primary bile acids (Zhao et al., 2013). In another work, the decreased abundance of *Erysipelotrichaceae* affected by antibiotics in human gut microbiota was reported by Yun (Yun et al., 2017). In addition to the altered richness of *Erysipelotrichaceae*, the reduced abundance of *Lactobacillaceae* and *Bacteroides* was also found to be in line with a review on the modification of gut microbiota by antibiotics (Ianiro et al., 2016). Notably, *Lactobacillaceae* is one of the beneficial bacterial species, the abundance of which could significantly decrease by the treatment of antibiotics particularly lincomycin (Tang et al., 2021), and the decrease was shown in the current study as well.

### Correlation of the changes in the microbiota with those in bile acid metabolism

The correlation of the effects of the antibiotics on the microbiota with their effects on the bile acid profiles may shed some light on the microbiota-related changes in bile acid profiles upon treatment with lincomycin or clindamycin, which is presented by the pearson correlation coefficients as shown in **Fig. 8**. *Erysipelotrichaceae* as well as *Akkermansiaceae* were shown highly relative to the bile acid profiles especially secondary bile acids involving DCA, LCA, ωMCA, and HDCA. In particular, *Erysipelotrichaceae* which showed a positive correlation with the levels of fecal secondary bile acids significantly reduced on the relative abundance in antibiotic-treated samples, leading to the decreased proportions of DCA, LCA, ωMCA, and HDCA (**Fig. 4**) after the 24 h incubation, whereas *Akkermansiaceae* presented the negative relation to the production of fecal secondary bile acids. Meanwhile, *Bacteroidaceae* also showed a significant positive correlation with the fecal secondary bile acids, and the reduced richness of *Bacteroidaceae* could result in the decreased production of secondary bile acids, especially in clindamycin-treated samples. In addition, *Prevotellaceae* and *Lactobacillaceae* were also shown to be highly relative to the secondary bile acids in the lincomycin- and clindamycin- treated groups respectively.

**Fig. 8.**
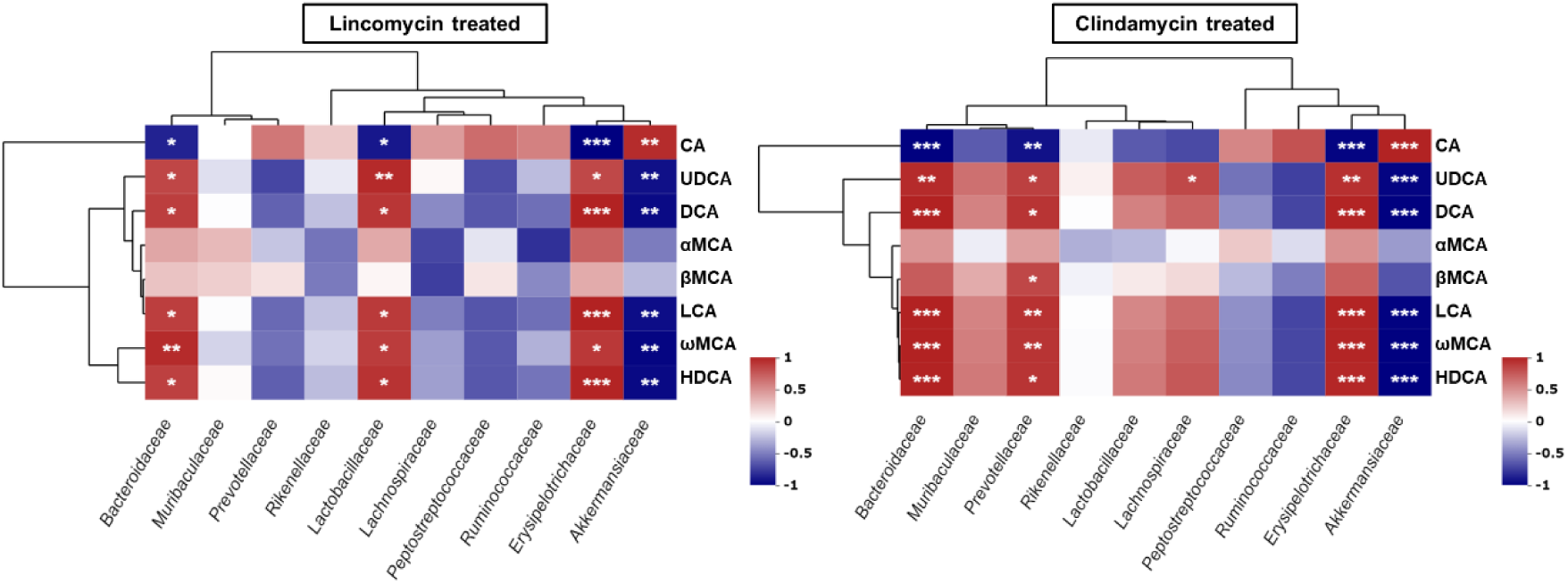
The correlation between bile acid and gut bacterial profiles. The heatmap presentation by the pearson correlation coefficients of the ten dominant families in rat fecal samples with the individual fecal bile acids including CA, UDCA, DCA, αMCA, βMCA, LCA, ωMCA, and HDCA is shown. All the fecal samples were taken at the 24 h of anaerobic incubation, and results are separately shown for lincomycin- and clindamycin- treated groups. The positive and negative correlation are presented by colours of red and blue respectively (*p<0.05, **p<0.01, ***p<0.001).

Together with the bile acid transformation pathways shown in **Fig. 6** and the previously reported studies, the correlation between the bacterial species and the conversion of individual bile acids could be more clear. For example, the accumulation of CA and the reduction in DCA may result from a reduction in 7α-dehydroxylation activity, which may be attributed to the loss of microorganisms belonging to the phylum *Firmicutes* known to contain 7α-dehydroxylation activity (Gérard, 2013; Jia et al., 2018), such as the reduced abundance of *Lactobacillaceae* and *Erysipelotrichaceae* observed in the current study. Meanwhile, as it is known that epimerization and oxidation of bile acids are also carried out by gut microbes (Guzior et al., 2021), the reduced relative abundance of microbes that possess epimerization and oxidation capabilities would result in decreased production of some secondary bile acids, such as ωMCA and LCA. As observed in this study, the reduced production of ωMCA and LCA upon antibiotic administration could be suspected to be associated with the weakened epimerization of βMCA and the weakened oxidation of CA (converting to 7,12-dioxo-LCA), respectively, and further related to the bacterial species of *Bacteroidaceae*, which has also been reported by Jia (Jia et. al., 2018). In addition to dehydroxylation, oxidation, and epimerization, deconjugation is another traditional distinct pathway related to microbial transformations of bile acids (Guzior et al., 2021). For example, the accumulation of CA could be caused by the weakening of deconjugation from TCA, and the deconjugation is associated with the effects on bile salt hydrolases (BSHs) linked to the microbial species belonging to *Bacteroidetes* and *Firmicutes* such as *Bacteroidaceae* and *Lactobacillaceae*, which was previously reported as well (Jia et. al., 2018).

The changes observed in the primary and secondary bile acids after lincosamide administration following the 24 h incubation revealed that antibiotic treatment had an influence on the gut microbiome, subsequently resulting in effects on bile acid metabolism and profiles, with effects detected in the newly applied *in vitro* model being comparable to what has been observed *in vivo*.

## 4. Conclusion

In this study, we demonstrate the utility of using an anaerobic fecal batch culture system to model the effects of antibiotics on bacterial composition and the resulting metabolism of individual primary and secondary bile acids using 16S rRNA gene sequencing and LC‒MS/MS. The consistent consequences for changes in fecal bile acid profiles as acquired from this *in vitro* model and a previous rat *in vivo* study, provide a proof of principle for the application of the developed 24 h *in vitro* fermentation batch model to study compound induced alterations in gut microbiota-dependent bile acid profiles. The method will facilitate elucidation of effects of other xenobiotics on the gut bacterial community and its consequences for bile acid metabolism, thereby contributing to the 3Rs (replacement, reduction and refinement) in animal testing.

## Acknowledgments

The authors are much appreciative of Jacques Vervoort for his supervision and efforts during the whole work. We are grateful to Sebastiaan Wesseling for help with the LC‒MS/MS work and Bert Spenkelink for help with arranging chemicals and collecting fecal samples.

We thank the funding supported by the Key Lab of Agro-product Quality and Safety (Beijing, China). The authors also acknowledge Ashwarya Murali and Ben van Ravenzwaaij from BASF (Ludwigshafen, Germany) for providing the rat fecal samples. IMCMR and MB would like to acknowledge the Long-range Research Initiative (LRI), which is an initiative from the European Chemical Industry Council, supporting the related LRI-ELUMICA project on the metabolic capacity of the intestinal microbiome.

## Authorship contribution statement

**Weijia Zheng**: Method development, Data curation, Formal analysis, Writing - original draft. **Wouter Bakker**: Method development. **Marta Baccaro**: Data curation. **Maojun Jin**: Supervision. **Jing Wang**: Supervision, Funding acquisition. **Ivonne M.C.M. Rietjens**: Supervision, Writing – review and editing.

## Conflict of interest

The authors declare that they have no conflicts of interest.

## Data availability

The data that has been used is confidential.

## Notes

### Competing Interest Statement

The authors have declared no competing interest.

